# Biomimetic, smart and multivalent ligands for G-quadruplex isolation and bioorthogonal imaging

**DOI:** 10.1101/2020.12.18.422878

**Authors:** Francesco Rota Sperti, Thibaut Charbonnier, Pauline Lejault, Joanna Zell, Claire Bernhard, Ibai E. Valverde, David Monchaud

**Affiliations:** ICMUB, CNRS UMR6302, UBFC Dijon, 9, Avenue Alain Savary, 21078 Dijon, France

## Abstract

G-quadruplexes (G4s) continue to gather wide attention in the field of chemical biology as their prevalence in the human genome and transcriptome strongly suggests that they may play key regulatory roles in cell biology. G4-specific, cell-permeable small molecules (G4-ligands) innovately permit the interrogation of cellular circuitries in order to assess to what extent G4s influence cell fate and functions. Here, we report on multivalent, biomimetic G4-ligands referred to as TASQs that enable both the isolation and visualization of G4s in human cells. Two biotinylated TASQs, **BioTASQ** and **BioCyTASQ**, are indeed efficient molecular tools to fish out G4s of mixtures of nucleic acids through simple affinity capture protocols and to image G4s in cells *via* a biotin/avidin pretargeted imaging system first applied here to G4s, found to be a reliable alternative to *in situ* click chemistry.

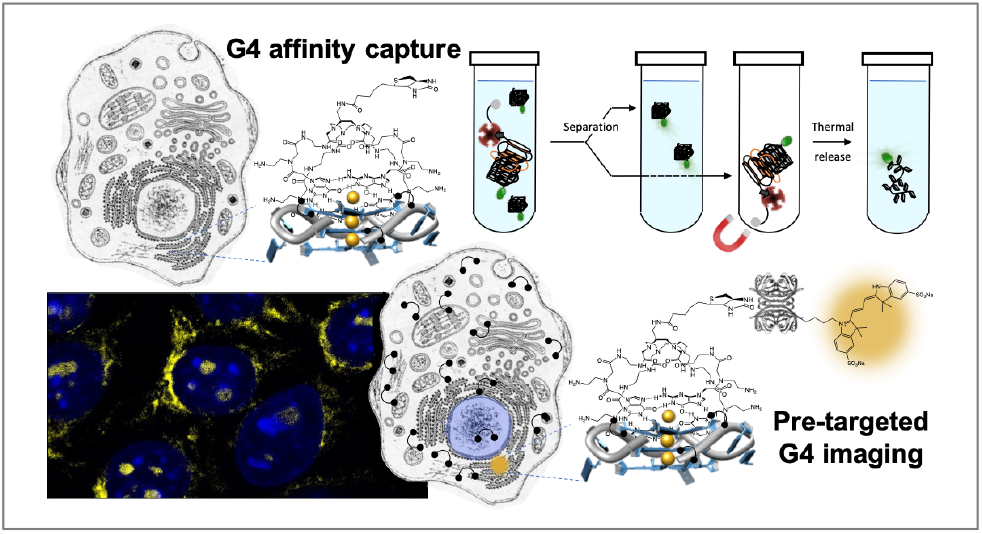

## Introduction

The principle of template-assembled synthetic proteins (TASP)^*1, 2*^ was developed by Mutter to tackle the complicated issue of how to control the folding of polypeptides into protein-like functional macromolecules. Inspired by this principle, Sherman reported on template-assembled synthetic G-quartets (TASQ)^*3*^ as a model system of how discrete G-quartets can be assembled intramolecularly. To this end, four guanine residues were covalently linked to a template resulting in a conformationally dynamic suprastructure, the guanines being either independent of each other (the so-called ‘open’ conformation of TASQ) or assembled into a G-quartet (*via* the formation of 8 hydrogen bonds, the ‘closed’ conformation).

The first prototypes of TASQ were built on a lipophilic template (Cram’s bowl-shape cavitands^*4*^ with long alkyl chains) for studying the cation (Na^+^, K^+^, Sr^2+^) chelation properties of discrete, synthetic G-quartets in organic media (CHCl_3_).^*3, 5, 6*^ Soon after, water-soluble TASQs were developed, hinging on hydrophilic templates such as the polyazamacrocyclic DOTA for the **DOTASQ**,^*7*^ the cyclodecapeptide RAFT (regioselectively addressable functionalized template, initially introduced by Mutter for the synthesis of TASP)^*2, 8, 9*^ for the **RAFT-G4**,^*10*^ as well as cavitands with phosphate appendages.^*11, 12*^ This opened new possibilities for biodirected applications such as the design of biomimetic G-quadruplex ligands (or G4-ligands,^*13*^ *vide infra*) or of molecular platforms to evaluate the G-quartet interacting properties of G4-ligands.^*12, 14*^

We have been particularly interested in fine-tuning the chemical scaffold of water-soluble TASQ in order to optimize their properties as biomimetic G4-ligands. The main binding site of a small-molecule within the G4 architecture is the external, accessible G-quartet.^*15*^ Since a G4 is more stable if it has more constitutive G-quartets,^*16*^ we set out to synthesize TASQ to interact with G4s according to such a biomimetic, like-likes-like interaction between a native G4 quartet and the synthetic TASQ quartet. By doing so, we have gradually modified the chemical nature of TASQ from the first biomimetic G4-ligand (**DOTASQ**,^*7, 17*^ and **PorphySQ**)^*18*^ to the first smart G4-ligands ^**PNA**^**DOTASQ**^*19, 20*^ (Figure 1) and ^**PNA**^**PorphySQ**,^*21*^ whose closed conformation is triggered only by interaction with G4s), twice-as-smart G4-ligands **PyroTASQ**^*22, 23*^ and **N-TASQ**,^*24, 25*^ smart ligands and smart probes, used to detect G4s *in vitro* and *in cella*, and the multivalent G4-ligand **BioTASQ**^*26, 27*^ (Figure 1), used as molecular bait for isolating G4 from human cells and identifying them by sequencing.

**Figure 1.**
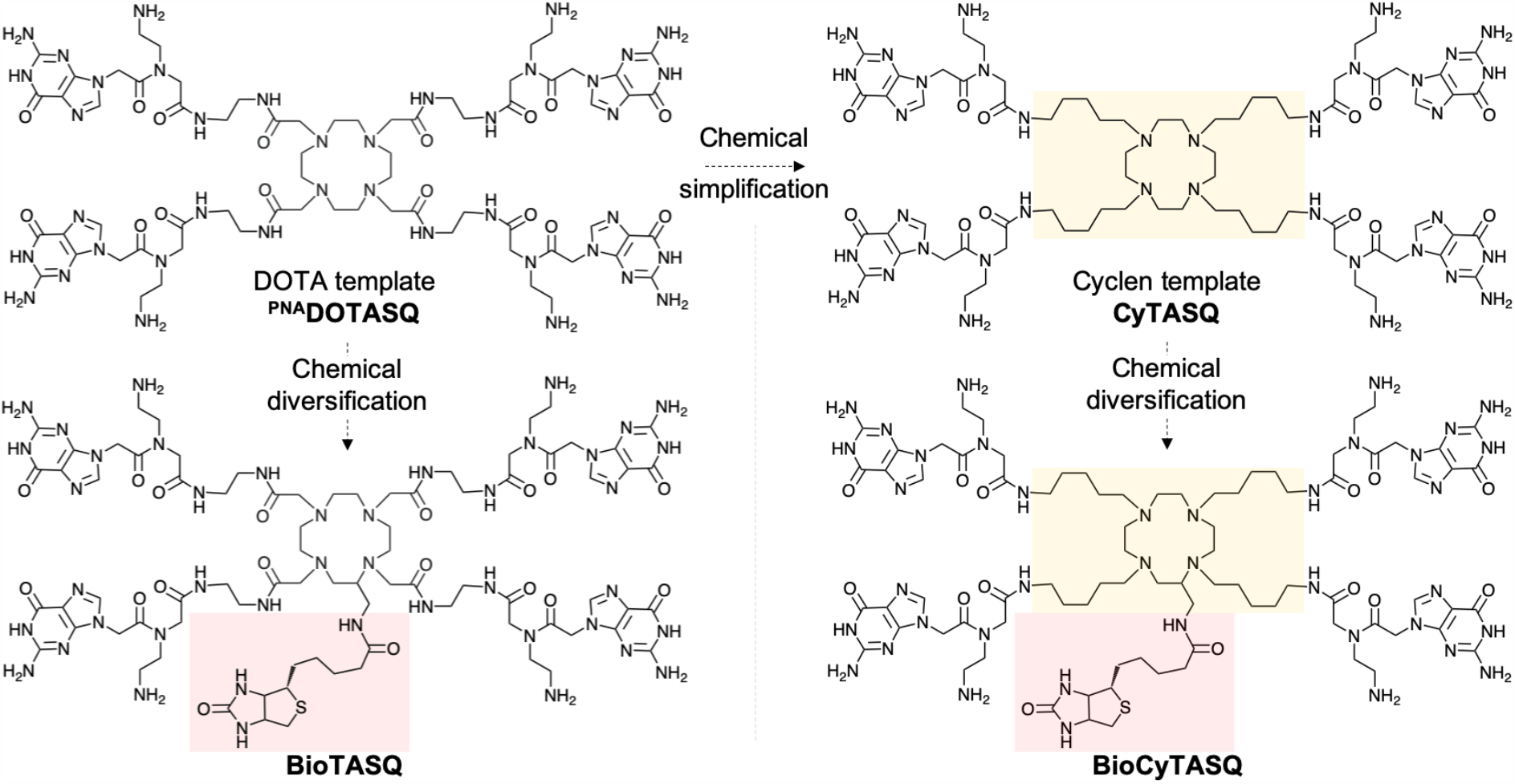
Structures of the DOTA-based ^PNA^DOTASQ and BioTASQ and the cyclen-based CyTASQ and BioCyTASQ.

A critical bottleneck in the development of TASQ resides in their chemical accessibility. Despite their relatively short synthesis, some technical pitfalls preclude efficient large-scale synthesis. We thus decided to revisit the synthesis of TASQ and report herein on the synthesis of both cyclen-template synthetic G-quartet (**CyTASQ**) and its biotinylated counterpart **BioCyTASQ** (Figure 1), whose design was inspired by that of the reference compound ^**PNA**^**DOTASQ** and **BioTASQ**,^*19, 26, 27*^ but with a simpler chemical scaffold and less time-consuming chemical accessibility.

### Design & synthesis of CyTASQ and BioCyTASQ

These two new TASQs were designed to keep the global structural organization of ^**PNA**^**DOTASQ** and **BioTASQ** (Figures 1 and 2), changing the topologically constrained amide linkers for more flexible alkyl linkers of similar length (in terms of number of atoms). This replacement, although conceptually simple, results in a more straightforward accessibility to the final TASQ with increased overall chemical yields (from 0.6% to 6% for **BioTASQ** and **BioCyTASQ**, respectively, Figure 2).

**Figure 2.**
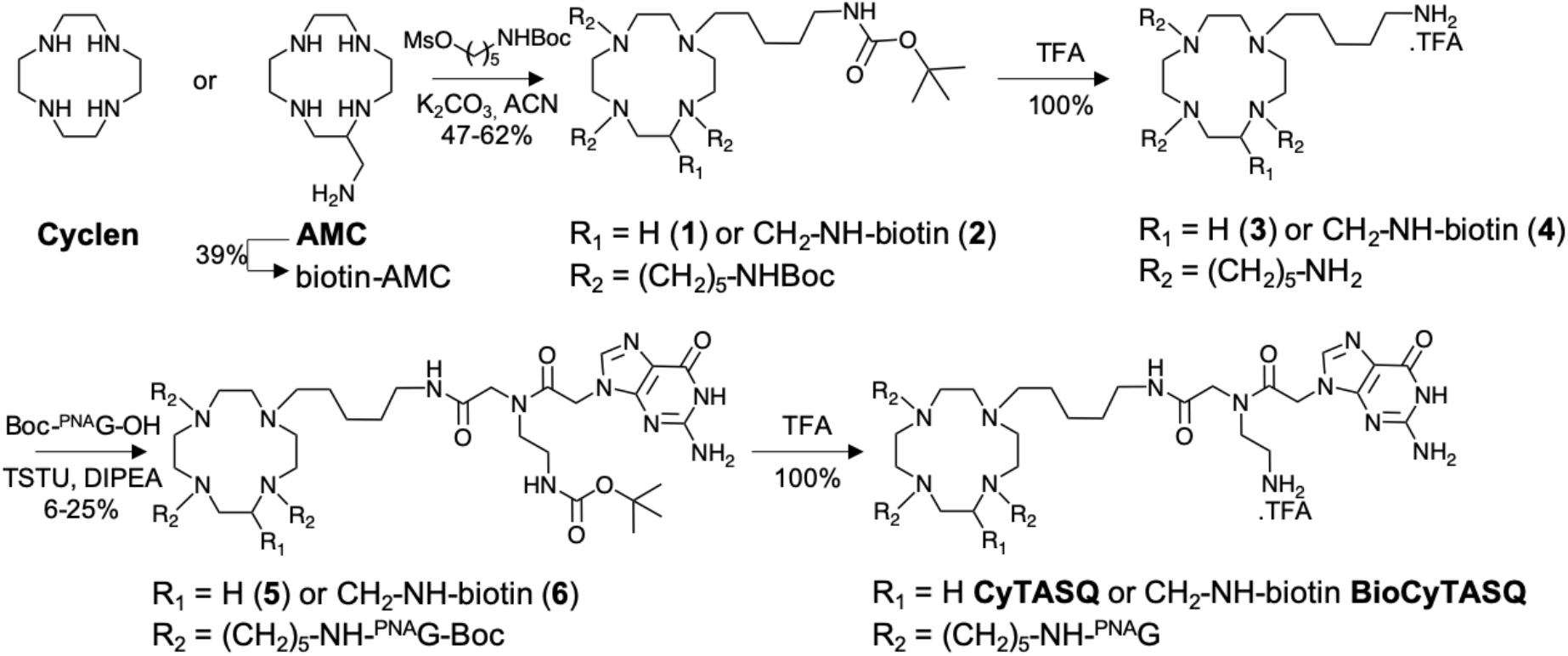
Chemical synthesis of CyTASQ and BioCyTASQ.

Briefly, the starting material for the synthesis of **CyTASQ** is the commercially available cyclen while that of **BioCyTASQ** is a biotinylated aminomethylcyclen (biotin-AMC),^*26, 27*^ prepared in one step (39% chemical yield) from AMC.^*28*^ These polyazamacrocycles were reacted with an excess (8 mol. equiv.) of 5-(Boc-amino)pentyl mesylate (prepared in two steps from the commercially available 5-amino-1-pentanol, 56% yield) in presence of potassium carbonate (K_2_CO_3_) in acetonitrile (ACN) to lead to compounds **1** (47% yield) and **2** (62% yield). These intermediates were deprotected with trifluoroactetic acid (TFA) to afford compounds **3** and **4** (100% yield), which were subsequently coupled with an excess (4.4 mol. equiv.) of Boc-^PNA^G-OH monomers (prepared in one step from the commercially available Boc-^PNA^G(Z)-OH, 72%) in presence of *N,N,Nʹ,Nʹ*-tetramethyl-O-(*N*-succinimidyl)uronium tetrafluoroborate (TSTU) and diisopropylethylamine (DIPEA) to afford the protected TASQs **5** (6% yield) and **6** (25% yield). The final compounds, **CyTASQ** and **BioCyTASQ**, were obtained after a last, quantitative deprotection step (TFA).

### Evaluation of G4-interacting properties of CyTASQ and BioCyTASQ *in vitro*

The apparent affinity of TASQs for different DNA/RNA sequences was evaluated *via* the firmly established FRET-melting assay (Figure 3A).^*29, 30*^ Doubly labelled biologically relevant G4-forming DNA/RNA sequences^*31*^ (0.2 μM) were heated from 20 to 90 °C in presence of TASQs (1.0 μM). The sequences used were the human telomeric-mimicking F-21-T (FAM-d[^5’^G_3_(T_2_AG_3_)_3_^3’^]-TAMRA; FAM for fluorescein amidite, TAMRA for tetramethylrhodamine), the Myc promoter-mimicking F-Myc-T (FAM-d[^5’^GAG_3_TG_4_AG_3_TG_4_A_2_G^3’^]-TAMRA), the human telomeric transcript F-TERRA-T (FAM-r[^5’^G_3_(U_2_AG_3_)_3_^3’^]-TAMRA) and the 5’-UTR of the mRNA coding for VEGF (F-VEGF-T, FAM-r[^5’^G_2_AG_2_AG_4_AG_2_AG_2_A^3’^]-TAMRA), along with F-duplex-T as a control (the hairpin-forming FAM-d[^5’^(TA)_2_GC(TA)_2_T_6_(TA)_2_GC(TA)_2_^3’^]-TAMRA).

**Figure 3.**
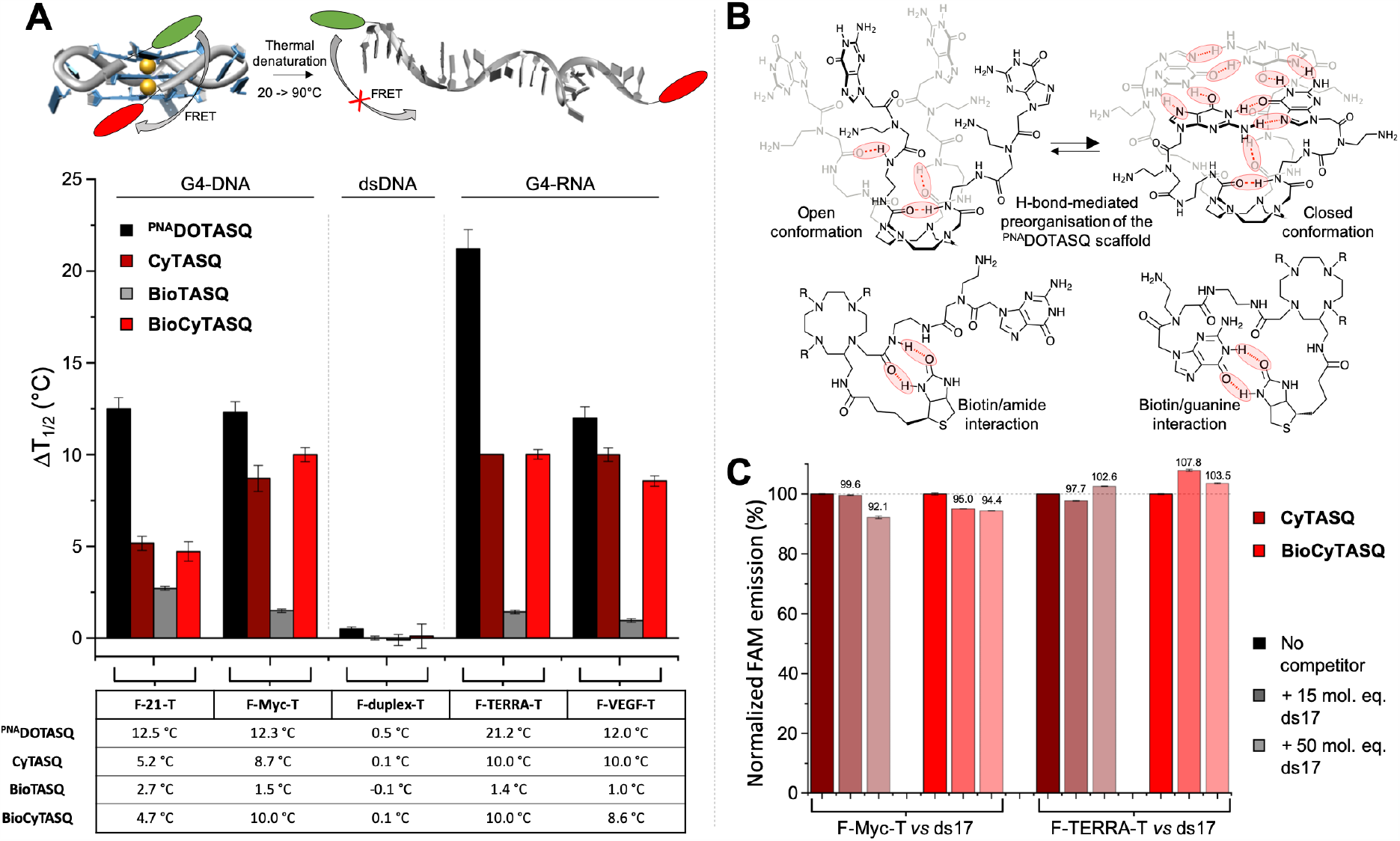
**A**. Schematic representation of the FRET-melting assay (upper panel) and results collected with doubly labelled G4s (DNA: F-21-T, F-Myc-T; RNA: F-VEGF-T, F-TERRA-T) and a duplex as a control (F-duplex-T) in the presence of TASQ (^PNA^DOTASQ in black, CyTASQ in brown, BioTASQ in gray and BioCyTASQ in red). **B**. Schematic representation of possible intramolecular H-bond between the biotin appendage and the amide linkage of both ^PNA^DOTASQ and BioTASQ arms. **C**. Competitive FRET-melting experiments performed with doubly labelled G4-DNA (F-Myc-T) or G4-RNA (F-TERRA-T) in presence of TASQ (CyTASQ in brown, BioCyTASQ in red) and increasing amounts of unlabelled duplex-DNA competitor (ds17, 15 and 50 molar equiv.).

As seen in figure 3A, the thermal stabilization of the DNA/RNA sequences was delayed in presence of all TASQs, but with a wide range of effects, with mid-transition temperatures (ΔT_1/2_, in °C) ranging from 1.0 and 21.8 °C. Interestingly, the chemical simplification of the G arms (**CyTASQ** *versus* ^**PNA**^**DOTASQ** and **BioCyTASQ** *versus* **BioTASQ**) affects the TASQ G4-stabilizing properties differently: on one hand, the stabilizations induced by ^**PNA**^**DOTASQ** (black bars, ΔT_1/2_ between 12.0 and 21.2 °C) are systematically higher than that of **CyTASQ** (brown bars, ΔT_1/2_ = 5.2 - 10.0 °C); whereas on the other hand, the **BioTASQ** stabilization values (gray bars, ΔT_1/2_ = 1.0 - 2.7 °C) are systematically lower than that of **BioCyTASQ** (red bars, ΔT_1/2_ = 4.7 - 10.0 °C), being comparable to that of **CyTASQ**. This could be interpreted in terms of flexibility of the G arms: arms with a greater flexibility may be detrimental for the non-biotinylated TASQs (**CyTASQ** *versus* ^**PNA**^**DOTASQ**), implying that the amide connection in the arms of ^**PNA**^**DOTASQ** somehow pre-organizes its external G-quartet for a better G4 recognition (presumably *via* internal H-bonding, Figure 3B).^*7*^ However, this flexibility improves the performances of biotinylated TASQs (**BioCyTASQ** *versus* **BioTASQ**) by decreasing the propensity of the biotin tag to intramolecularly interact with one of the G arms of the TASQ, known to perturb their association with G4 and initially thought to be a biotin/guanine interaction.^*27*^ These new results suggest that the interaction might occur between the biotin and the amide connector of the G arm neighboring the cyclen (Figure 3B), which seems to be less sterically constrained. **CyTASQ** and **BioCyTASQ** are thus less structurally preorganized than ^**PNA**^**DOTASQ** (lower apparent affinity) but also less internally poisoned than **BioTASQ** (higher apparent affinity).

These results also show that TASQs do not interact with duplex-DNA (see the control with F-duplex-T, with ΔT_1/2_ < 0.5 °C), highlighting a G4 specificity that was confirmed by competitive FRET-melting assays performed with F-Myc-T and F-TERRA-T (0.2 μM) in presence of **Bio**(**Cy**)**TASQ** (1.0 μM) and an excess (3 and 10 μM) of an unlabelled duplex-DNA competitor ds17 (d[^5’^C_2_AGT_2_CGTAGTA_2_C_3_^3’^]/d[^5’^G_3_T_2_ACTACGA_2_CTG_2_^3’^]). The new TASQs are highly G4-specific (Figure 3C) with a maintained stabilization of >92% for **CyTASQ** and >98% for **BioCyTASQ**. The high affinity and exquisite selectivity of the biotinylated **BioCyTASQ** makes it well suited to be used as a molecular bait for fishing G4s out of nucleic acids mixtures.

### Evaluation of G4-capture properties of CyTASQ and BioCyTASQ *in vitro*

The ability of the biotinylated TASQs to pull down G4s was assessed *via* an optimized version of the ‘pull-down’ or ‘affinity capture’ protocol developed for the **BioTASQ** (Figure 4A).^*27*^ FAM-labelled G4-forming DNA/RNA sequences (1 μM) were incubated for 2 h with either **BioTASQ** or **BioCyTASQ** (10 μM) in presence of streptavidin-coated magnetic beads in TrisHCl buffer (20mM, pH 7.2) containing 1 mM KCl, 99 mM LiCl and 10 mM MgCl_2_ (the Mg^2+^ being found to decrease the random electrostatic interactions between DNA and TASQs). The G4-forming sequences used were the human telomeric-mimicking F-22AG (FAM-d[^5’^AG_3_(T_2_AG_3_)_3_^3’^]), a section of the promoters of MYC (FAM-d[^5’^GAG_3_TG_4_AG_3_TG_4_A_2_G^3’^]) and SRC genes (FAM-d[^5’^G_3_AG_3_AG_3_CTG_5_^3’^]), and F-duplex as a control (FAM-d[^5’^(TA)_2_GC(TA)_2_T_6_(TA)_2_GC(TA)_2_^3’^]). The FAM-G4/**Bio**(**Cy**)**TASQ**/beads assemblies were isolated (magnetic immobilization), the supernatant removed and the FAM-G4 resuspended in solution after a thermal denaturation step (10 min at 90 °C). The capture/release efficiency of TASQ was quantified by the FAM emission of the resulting solution, normalized to the controls performed without TASQ.

**Figure 4.**
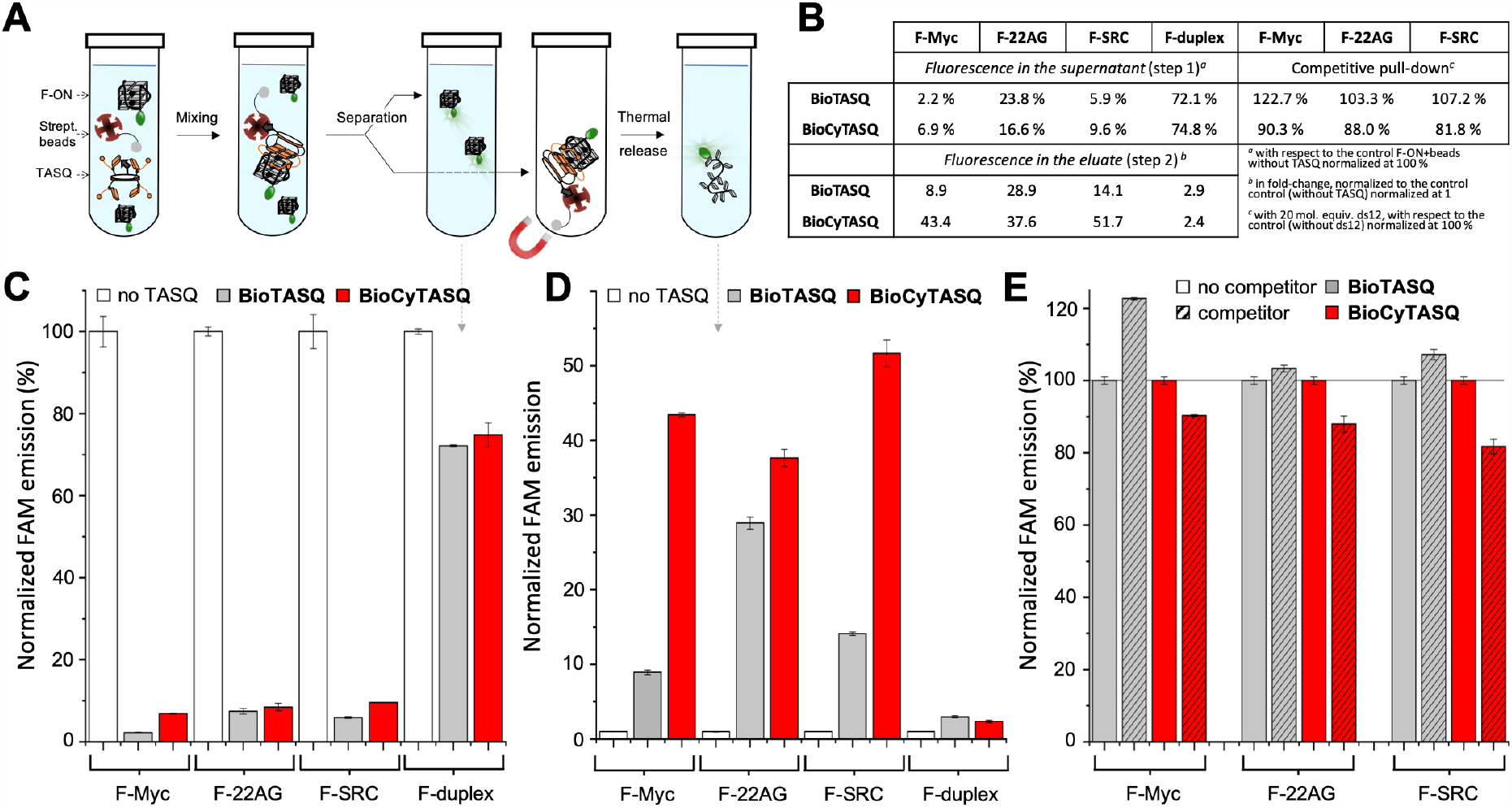
**A**. Schematic representation (**A**) and results (**B**) of the *in vitro* G4 pull-down protocol performed with FAM-labelled oligonucleotides (the G4s F-Myc, F-22AG and F-SRC; the hairpin F-duplex) and either **BioTASQ** or **BioCyTASQ**, quantified by either the decrease of the fluorescence of the supernatant (step 1, **C**) or the increase of the fluorescence during the elution (step 2, **D**). Competitive pull-down experiments (**E**) performed with F-Myc, F-22AG and F-SRC, **Bio**(**Cy**)**TASQ** and the unlabelled duplex competitor ds12 (20 mol. equiv.).

Both **BioTASQ** and **BioCyTASQ** efficiently captured G4s in solution (Figures 4B-E). This could be assessed by both the decrease of the FAM intensity of the supernatant, which reflects the pull-down *per se* (with a pull-down efficiency as low as -98 % as compared to the control, *i*.*e*., experiments performed without TASQ, normalized to 100%, panel C) and the increase of the FAM intensity after releasing the beads content by a thermal denaturation step (with a release enrichment up to 52-fold as compared to the control, *i*.*e*., experiments performed without TASQ, normalized to 1, panel D). The results obtained again demonstrate the improved performances of **BioCyTASQ** (red bars) in comparison with **BioTASQ** (gray bars) during the elution step (9- *versus* 44-, 29- *versus* 38- and 14- *versus* 52-fold enrichment for F-Myc, F-22AG and F-SRC, respectively). Again, these results also show that TASQs are specific for G4s (see the control with F-duplex, with 2- and 3-fold enrichment), a specificity further confirmed by competitive pull-down experiments performed with F-Myc, F-22AG and F-SRC (1 μM) in presence of **Bio**(**Cy**)**CyTASQ** (10 μM) and an excess (20 μM) of an unlabelled duplex-DNA competitor ds12 (the self-complementary strands d[^5’^CGCGA_2_T_2_CGCG^3’^]), in which the pull-down efficiency is maintained at >82% for **BioTASQ** and >103% for **BioCyTASQ** (Figure 4E).

### Evaluation of G4-interacting properties of CyTASQ and BioCyTASQ *in cella*

The biotin appendage of **Bio**(**Cy**)**TASQ** can also be used for optical imaging. Inspired by pretargeted imaging and therapy strategies involving biotin- and/or streptavidin-antibody conjugates,^*32-34*^ we used **Bio**(**Cy**)**TASQ** for imaging G4s in human cancer cells (MCF7) based on the highly specific interaction between the biotin appendage of TASQ and a Cy3-labelled streptavidin (SA-Cy3). Two strategies were implemented, with a systematic comparison of the properties of **BioTASQ** and **BioCyTASQ**: a post-fixation or a live-cell labelling strategy. In the first strategy, MCF7 cells are fixed (MeOH) then incubated sequentially with **Bio**(**Cy**)**TASQ** (1 μM, 1 h), SA-Cy3 (1 μg/mL) and DAPI (2.5 μg/mL) (Figure 5A). In the second strategy, MCF7 cells are treated live with **Bio**(**Cy**)**TASQ** (1 μM, 24 h; IC_50_ >100 μM) before fixation (MeOH), and sequential SA-Cy3 and DAPI labelling (Figure 5B). Both strategies are intended to provide complementary outputs as the former leads to a snapshot of G4 landscapes as captured by chemical fixation while the latter highlights cellular sites where TASQs accumulate in live cells.

**Figure 5.**
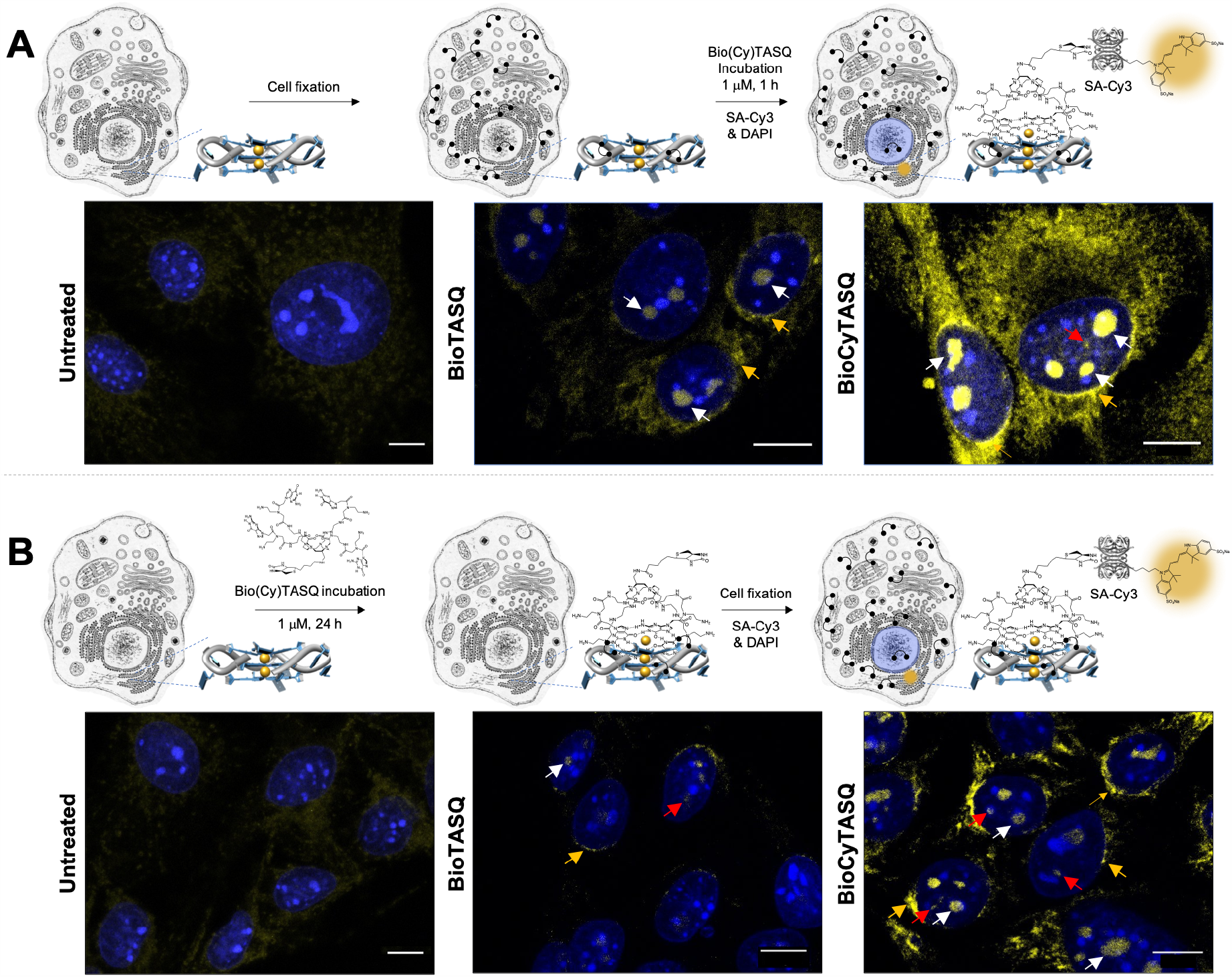
Schematic representation (and images) of the **Bio**(**Cy**)**TASQ**-based, two-step strategies implemented to label G4s in MCF7 cells, either (**A**) post-fixation labelling or (**B**) live-cell incubation. Images are collected in the range of 518-562 nm for SA-Cy3 (yellow) and < 425 nm for DAPI (blue channel); scale bar 10 μm; yellow, white and red arrows indicate G4 *foci* found in perinuclear regions, nucleoli and nucleoplasm, respectively.

In both instances, **BioCyTASQ** provided a brighter response than **BioTASQ** (Figure 5, the images were collected under strictly identical experimental setups). Post-fixation labelling led to a rather diffuse labelling in the cytoplasm along with stronger labelling in perinuclear regions (yellow arrows), nucleoli (white arrows) and non-nucleoli nucleoplasmic sites (red arrows). Live-cell labelling provided a similar perinuclear/nucleoli/nuclear distribution (yellow/white/red arrows) but with higher contrast, highlighting G4 *foci* in a more precise manner, with abundant G4-RNA sites, mostly found in ribosome sites (nucleoli (white arrows) and rough endoplasmic reticulum (yellow arrows)) and sparser G4-DNA sites (located in the nucleoplasm (red arrows)). These results are fully in line with the direct G4 labelling pattern provided by N-TASQ.^*24, 25, 35, 36*^ We observe that the live-cell treatment shows a similar staining pattern to post-fixation labelling, indicating the G4 landscape is not redistributed by TASQ treatment. Live cell imaging is more accurate because the ligand’s target is not chemically altered by fixation. This approach should thus be preferred to investigate G4 biology with TASQs. To the best of our knowledge, this is the first description of a pretargeted imaging strategy based on the biotin/avidin system applied to visualize G4s in human cells.

## Conclusion

The panel of strategies now available for gaining insights into G4 landscapes within human cells encompasses optical imaging^*37, 38*^ and sequencing methods.^*39, 40*^ These strategies rely on exquisitely efficient and specific molecular tools to either label G4 sites within cells (*e*.*g*., direct labelling with G4 probes or indirect labelling *via* bioorthogonal chemistry)^*37, 38, 41, 42*^ or stabilize G4s *in vitro* to cause polymerase stalling which is used as a next-generation sequencing readout (*e*.*g*., pyridostatin in G4-seq^*43*^ and rG4-seq^*44*^ protocols).^*39, 40, 45*^ In light of the number of potential G4-forming sequences in the human genome and transcriptome,^*43, 46-52*^ and the biological processes they might be involved in,^*31, 53-56*^ the task can seem daunting; however, this offers a virtually unlimited playground for chemical biology investigation, as every new tool and approach could provide alternative and/or complementary information helping to further decipher the complex biology of G4s.

Here, we continue to exploit the potential of biomimetic and smart G4 ligands, demonstrating the versatility of biotinylated TASQs to both isolate G4s by pull-down affinity capture, implementable in cells by G4RP-seq for instance, and visualize G4s *via* an original streptavidin/biotin pretargeted imaging approach, found to be a reliable alternative to *in situ* click chemistry.^*42*^ We address the problem of poor chemical accessibility of TASQs *via* the design and synthesis of the new generation TASQs rereferred to as **CyTASQ** and **BioCyTASQ**, which can now be produced on scales and timescales compatible with a broader use. The *in vitro* validation provided here lends further credence to the strategic relevance of TASQs as multivalent molecular tools to investigate G4s *in cella* endowing them with new functionalities that continue to demonstrate the versatility of their applications.

## Methods

### Synthesis of CyTASQ

Compound **1**: To a solution of cyclen (30 mg, 0.20 mmol, 1.0 equiv.) in acetonitrile (1.16 mL) was added 5-((tert-butoxycarbonyl)amino)pentyl methanesulfonate (400 mg, 1.40 mmol, 7.0 equiv.) and K_2_CO_3_ (191 mg, 1.40 mmol, 7.0 equiv.) and the solution was stirred until complete conversion of the starting material (48h, monitoring by RP-HPLC). The crude mixture was filtered, and concentrated under vacuum. The residue was then purified by silica gel column chromatography (CH_2_Cl_2_/MeOH (90:10)). After evaporation of the solvents, compound **1** was obtained (74.2 mg, 0.08 mmol, 47 % yield). ^1^H NMR (500 MHz, CDCl_3_): δ 4.81 (br s, 4H), 3.10 (dt, *J* = 6.3, 6.3 Hz, 8H), 2.89-2.73 (m, 16H), 2.08 (s, 3H), 1.54-1.49 (m, 17H), 1.43 (s, 36H), 1.31 (m, 9H). ESI-MS: [M+H]^+^ m/z = 913.95 (calcd. for C_48_H_97_N_8_O_8_: 913.74). Compound **3**: compound **1** was stirred in 2 mL of TFA for 1 hour. After evaporation of the TFA, compound **3** was obtained (78.5 mg, 0.08 mmol, 100 % yield). Compound **5**: Boc-^PNA^G-OH (160 mg, 0.40 mmol, 4.5 equiv.) and TSTU (118 mg, 0.40 mmol, 4.5 equiv.) were dissolved in DMF (1 mL) and DIPEA was added (54 μL, 0.36 mmol, 4.0 equiv.). After 1 h, a solution of compound **3** (45 mg, 0.09 mmol, 1.0 equiv.) and DIPEA (54 μL, 0.36 mol, 4.0 equiv.) in DMF (1 mL) was added and the mixture was stirred at RT for 3 d. The solution was then concentrated under vacuum and purified by RP-HPLC in a H_2_O/ACN + 0.1% TFA mixture (gradient of 5 to 40 % over 20 min). After evaporation of the solvents, the compound **5** was obtained (12.3 mg, 0.01 mmol, 6 % yield). ^1^H NMR (500 MHz, DMSO-d6) δ 10.94 (s, 4H), 8.31 (m, 1.3H), 8.14 (m, 0.7H), 8.02-7.78 (m, 7H), 7.03-7.01 (m, 2H), 6.76-6.55 (m, 9H), 5.08-5.00 (m, 5H), 4.88-4.84 (m, 3H), 4.19 (br s, 0.5H), 4.12 (br s, 2.5H), 3.97 (br s, 0.5H), 3.92 (br s, 0.5H), 3.87 (m, 4.5H), 3.72 (br s, 0.5H), 3.54-3.7 (m, 5H), 3.29-2.95 (m, 34H), 2.86-2.71 (m, 8H), 1.51-1.36 (m, 50H), 1.27-1.14 (m, 10H). ESI-MS: [M+H]^+^ m/z = 2079.17259 (calcd. for C_92_H_149_N_36_O_20_: 2079.17434). **CyTASQ**: compound **5** was dissolved in 1 mL of TFA and stirred for 1 hour. The complete deprotection was assessed by HPLC-MS (H_2_O/ACN + 0.1% TFA mixture (gradient of 5 to 100 % over 7 min); retention time = 0.44 min; [M+2H]^2+^ m/z = 840.0; calcd. for: C_72_H_117_N_36_O_12_ [M+2H]^2+^ m/z = 839.9). TFA was removed under reduced pressure, the residue lyophilized and used immediately, without further purification.

### Synthesis of BioCyTASQ

Compound **2**: To a solution of biotin (514.7 mg, 2.10 mmol, 0.8 equiv.) and DIPEA (949 μL, 5.40 mmol, 2.0 equiv.) in DMF (10 mL) was added TSTU (900 mg, 3.00 mmol, 1.1 equiv.) and the solution was stirred at RT until the complete activation of the carboxylic acid, assessed *via* HPLC-MS monitoring. The solution was then added dropwise (1 mL/3 h) to a solution of AMC (546 mg, 2.70 mmol, 1.0 equiv.) in DMF (20 mL) and the reaction was carefully monitored by HPLC-MS. Upon completion, TFA (500 μL) was added and the solution was concentrated under vacuum and the resulting residue purified by RP-HPLC in a H_2_O/can + 0.1% TFA mixture (gradient of 2 to 100 % over 50 minutes). After evaporation of the solvents, compound **2** was obtained (926 mg, 1.04 mmol, 39% yield). ^1^H NMR: (500 MHz, D_2_O) δ 4.60 (dd, *J* = 7.9, 4.9 Hz, 1H), 4.41 (dd, J = 8.0, 4.5 Hz, 1H), 3.51 – 2.73 (m, 20H), 2.30 (t, *J* = 7.3 Hz, 2H), 1.78 – 1.50 (m, 4H), 1.48 – 1.29 (m, 4H) (signals of intracyclic amines and amide are missing due to proton solvent exchange). ^13^C NMR: (126 MHz, D_2_O) δ 177.7, 165.4, 117.4, 115.1, 62.1, 60.3, 55.4, 51.9, 46.3, 44.3, 44.2, 44.0, 42.6, 42.1, 39.6, 39.1, 39.1, 35.3, 28.0, 27.9, 27.7, 24.8, 16.2. ESI-MS: [M+H]^+^ m/z = 428.3 (calcd. for C_19_H_38_N_7_O_2_S: 428.6). Compound **4**: To a solution of compound **2** (283.2 mg, 0.32 mmol, 1.0 equiv.) in acetonitrile (4 mL) was added 5-((tert-butoxycarbonyl)amino)pentyl methanesulfonate (720 mg, 2.56 mmol, 8.0 equiv.) and K_2_CO_3_ (472 mg, 3.42 mmol, 10.0 equiv.) and the solution was stirred at 50°C for 56 h. Another aliquot of 5-((tert-butoxycarbonyl)amino)pentyl methanesulfonate (392 mg, 1.40 mmol, 4.0 equiv.) was added and further stirred for 16 h. The crude mixture was filtered and concentrated under vacuum and the residue was purified by HPLC in a H_2_O/ACN + 0.1% TFA mixture (gradient of 30 to 100 % over 40 min). Compound **4** was obtained (232.5 mg, 0,20 mmol, 62% yield). ^1^H NMR: (500 MHz, DMSO-d6) δ 8.01 (s, 1H), 7.84 (s, 2H), 7.70 (s, 1H), 6.80 (d, *J* = 14.5 Hz, 3H), 6.40 (s, 2H), 4.32 (dd, *J* = 7.8, 4.9 Hz, 1H), 4.21 – 4.14 (m, 1H), 4.13 (dd, *J* = 7.8, 4.9Hz, 1H), 3.40 – 3.34 (m, 5H), 3.16 (d, *J* = 7.5 Hz, 2H), 3.09 (dt, *J* = 10.0, 5.3 Hz, 1H), 3.05 (s, 13H), 2.92 (tt, *J* = 10.6, 5.6 Hz, 8H), 2.80 (tt, *J* = 13.1, 5.7 Hz, 3H), 2.71 (s, 4H), 2.13 – 2.06 (m, 2H), 1.66 (tt, *J* = 14.1, 7.0 Hz, 2H), 1.54 (s, 11H), 1.53 (t, *J* = 11.4 Hz, 1H), 1.37 (d, *J* = 2.2 Hz, 39H), 1.30 (s, 9H), 1.28 (d, *J* = 7.8 Hz, 2H). ^13^C NMR: (126 MHz, DMSO-d6) δ 162.9, 158.5, 158.2, 155.8, 117.5, 115.1, 77.5, 70.6, 70.3, 61.2, 59.4, 55.6, 36.7, 30.8, 29.1, 28.4, 28.3, 22.4. MALDI: [M+H]^+^ 1168.09 m/z = (calcd. for C_59_H_114_N_11_O_10_S: 1168.68). Compound **6**: Boc-^PNA^G-OH (245 mg, 0.63 mmol, 4.8 equiv.), TSTU (191 mg, 0.63 mmol, 4.8 equiv.) were dissolved in DMF (1.5 mL) and DIPEA was added (99 μL, 0.63 mmol, 4.8 equiv.). After the complete activation of the carboxylic acid (HPLC-MS monitoring), a solution of compound **5** (100 mg, 0.13 mmol, 1.0 equiv.) and DIPEA (44 μL, 0.26 mmol, 2.0 equiv.) in DMF (1.5 mL) was added. The mixture was stirred at RT overnight. The solution was then concentrated under vacuum and the residue was purified by RP-HPLC in a H_2_O/ACN + 0.1% TFA mixture (gradient of 15 to 65 % over 50 minutes). After evaporation of the solvents, compound **6** was obtained (77.8 mg, 0.03 mmol, 25% yield). ^1^H NMR: (500 MHz, DMSO-d6) δ 10.8 (m, 2H), 8.3-7.9 (m, 7H),7.70 (s, 3H), 7.1 (s, 2H), 6.8 (s, 1H), 6.5 (m, 5H), 6.4 (m, 2H),5.0 (s, 4H),4.9 (m, 2H), 4.4 (m, 4H), 3.9-3.8 (m, 32H), 3.5 (m, 8H), 3.3 (m, 10H), 3.2 (m, 5H), 3.1 (m, 13H), 2.9 (m, 2H), 2.6 (m, 2H), 2.1 (m, 3H), 1.6 (m, 11H), 1.5 (s, 36H), 1.3 (m, 13H). ESI-HRMS: [M+H+Na]^2+^ m/z = 1178.63633 (calc. for [C_103_H_166_N_39_O_22_SNa]^2+^: = 1178.63519). **BioCyTASQ**: compound **6** was dissolved in 1 mL of TFA and stirred for 1 hour. The complete deprotection was assessed by HPLC-MS (H_2_O/ACN + 0.1% TFA mixture (gradient of 5 to 100 % over 7 min); retention time = 0.38 min; [M+2H]^2+^ m/z = 967.5; calcd. for C_83_H_134_N_39_O_14_S: [M+2H]^2+^ m/z = 967.6). TFA was removed under reduced pressure, the residue lyophilized and used immediately, without further purification.

### FRET-melting assays

FRET-melting experiments were performed in a 96-well format using a Mx3005P qPCR machine (Agilent) equipped with FAM filters (λ_ex_ = 492 nm; λ_em_ = 516 nm) in 100 μL (final volume) of 10 mM lithium cacodylate buffer (pH 7.2) plus 10 mM KCl/90 mM LiCl (F21T, F-duplex-T) or plus 1 mM KCl/99 mM LiCl (F-Myc-T, F-Terra-T, F-VEGF-T) with 0.2 μM of labeled oligonucleotide and 1 μM of TASQ. Competitive experiments were carried out with labeled oligonucleotide (0.2 μM), 1 μM TASQ and increasing amounts (0, 15 and 50 equiv.) of the unlabeled competitor ds17. After an initial equilibration step (25°C, 30 s), a stepwise increase of 1°C every 30s for 65 cycles to reach 90°C was performed, and measurements were made after each cycle. Final data were analyzed with Excel (Microsoft Corp.) and OriginPro^®^9.1 (OriginLab Corp.). The emission of FAM was normalized (0 to 1), and T_1/2_ was defined as the temperature for which the normalized emission is 0.5; ΔT_1/2_ values are means of 3 experiments.

### Pull-down assay

The streptavidin MagneSphere^®^ beads (Promega) were washed 3 times with TrisHCl buffer containing 1 mM KCl, 99 mM LiCl and 10 mM MgCl_2_. **Bio**(**Cy**)**TASQ** (10 μM) was mixed with 5’-labeled oligonucleotides (F-ON, 1 μM), *i*.*e*., F-Myc, F-SRC, F-22AG and F-duplex, MagneSphere^®^ beads (32 μg) in the same TrisHCl buffer (320 μL final volume) and stirred for 2 h at 25 °C. The beads were immobilized (magnet) and the supernatant removed (*fluorescence analysis 1*). The solid residue was resuspended in 320 μL of TBS 1X buffer, heated for 10 min at 90 °C (gentle stirring 800 r.p.m.) and then centrifuged for 2 min at 8900 rpm. The supernatant was taken up for analysis (magnet immobilization), after being distributed in 3 wells (100 μL each) of a 96-well plate, using a ClarioStar^®^ machine (BMG Labtech) equipped with FAM filters (λex = 492 nm; λem = 516 nm) (*fluorescence analysis 2*). Competitive experiments were carried out with labeled oligonucleotide (1 μM), 10 μM TASQ and in the presence of the unlabeled competitor ds12 (20 mol. equiv.). Data were analyzed with Excel (Microsoft Corp.) and OriginPro^®^9.1 (OriginLab Corp.); normalized FAM emission values are means of 3 measurements, according to the following methodology: *i*-*fluorescence analysis 1*: each analysis originated in 3 different experiments, performed as triplicates: a/ 3 control wells in which the F-ON was alone, whose FAM emission was normalized to 100; b/ 3 wells in which F-ON was mixed with beads, in order to quantify the non-specific F-ON/bead binding; and c/ 3 wells in which F-ON and **Bio**(**Cy**)**TASQ** were mixed with beads, in order to quantify the specific F-ON/**Bio**(**Cy**)**TASQ**/bead binding. *ii-fluorescence analysis 2*: each analysis originated in 2 different experiments, performed as triplicates: a/ 3 control wells comprising solutions that resulted from experiments performed with F-ON and beads, in order to quantify the non-specific F-ON/bead binding, the FAM emission of the solution was normalized to 1; and b/ 3 wells comprising solutions that resulted from experiments performed with F-ON, **Bio**(**Cy**)**TASQ** and beads, in order to quantify the actual **Bio**(**Cy**)**TASQ** capture capability when compared to the control experiments.

### Cell culture and imaging

MCF7 cells were routinely cultured in 75 cm^2^ tissue culture flasks (Nunc) at 37 °C in a humidified, 5% CO2 atmosphere in Dulbecco’s Modified Eagle Medium (DMEM) supplemented with 10% fetal bovine serum (FBS, Gibco) and 1% Penicillin-Streptomycin (Pen-Strep: 5.0 U.mL^−1^ Pen/5.0 μg.mL^−1^ Strep, Gibco) mixture. Cells were subcultured twice a week using standard protocols. Round coverslips (12 mm) were sterilised with 70% ethanol before cell seeding. MCF7 cells were seeded at a density of 6.10^4^ cells per coverslip on chambered coverslips (24 well-plate) and allowed to recover for 24 h. In the case of live-cell labelling, cells were incubated with **Bio**(**Cy**)**TASQ** (1 µM, 24h) at 37 °C, washed once with PBS 1X and fixed and permeabilized with ice cold MeOH for 10 min at room temperature. In the case of post-fixation labeling, seeded cells were fixed and washed with PBS 1X (3x), then incubated with 1 µM **Bio**(**Cy**)**TASQ** for 1 h at 25°C, washed with PBS 1X (3 × 5 min), then incubated for 1 h at 25°C in a light-tight box with Streptavidin-Cy3 (1 μg/mL) and washed with PBS 1X (3 × 5 min) and once with H_2_O. Cells were mounted onto glass microscope slides with Fluoromount-G (Southern Biotech) containing DAPI (2.5 μg/mL). The cells were imaged with a Leica TCS SP8 confocal laser-scanning microscope with a 63X oil objective, collected through the following channels: DAPI (excitation: 340-380 nm; emission: < 425 nm), GFP (excitation: 450-490 nm; emission: 500-550 nm) and Cy3 (excitation: 518-562 nm; emission: > 580 nm). Images were processed with ImageJ (https://fiji.sc).^*57*^

## Acknowledgments

The authors thank the Centre National de la Recherche Scientifique (CNRS), the Agence Nationale de la Recherche (ANR-17-CE17-0010-01; ANR-18-CE07-0017-03), the Université de Bourgogne, Conseil Régional de Bourgogne and the European Union (PO FEDER-FSE Bourgogne 2014/2020 programs) and the INSERM Plan Cancer 2014-2019 (n° 19CP117-00) for financial support. Souheila Amor and Katerina Duskova are warmly thanked for their help in the pretargeted imaging protocols; Christine Arnould (INARe Dijon) and DImaCell (*Dispositif Inter-régional d’Imagerie Cellulaire*) for the help with and access to the microscopes.

## Notes

### Competing Interest Statement

The authors have declared no competing interest.

